# Close kin dyads indicate intergenerational dispersal and barriers

**DOI:** 10.1101/2022.01.18.476819

**Authors:** Thomas L Schmidt, Samia Elfekih, Li-Jun Cao, Shu-Jun Wei, Mohamed B Al-Fageeh, Majed Nassar, Abdulaziz Al-Malik, Ary A Hoffmann

## Abstract

The movement of individuals through continuous space is typically constrained by dispersal ability and dispersal barriers. A range of approaches have been developed to investigate these. Kindisperse is a new approach that infers intergenerational dispersal (σ) from close kin dyads, and appears particularly useful for investigating taxa that are difficult to observe individually. This study, focusing on the mosquito *Aedes aegypti*, shows how the same close kin data can also be used for barrier detection. We empirically demonstrate this new extension of the method using genome-wide sequence data from 266 *Ae. aegypti*. First, we use the spatial distribution of full-sib dyads collected within one generation to infer past movements of ovipositing female mosquitoes. These dyads indicated the relative barrier strengths of two roads, and performed favourably against alternative genetic methods for detecting barriers. The difference in variance between the sib and first cousin spatial distributions was used to infer movement over the past two generations, providing estimates of intergenerational dispersal (σ = 81.5-197.1 m.gen^-1/2^) and density (ρ = 833-4864 km^-2^). Dispersal estimates showed general agreement with those from mark-release-recapture studies. Barriers, σ, ρ, and neighbourhood size (331-526) can inform forthcoming releases of dengue-suppressing *Wolbachia* bacteria into this mosquito population.

## Introduction

Dispersal is a central and widely studied process in ecology and evolution. Dispersal describes how organisms and their progeny come to occupy different locations, and it can refer to the movement of individuals or gametes (Howard 1960; Berry et al. 2004; Royle and Young 2008) or to the intergenerational outcomes of this movement (Wright 1943; Rousset 2000). Approaches for assessing dispersal at the individual level include those that measure movement rates over key life history stages, such as through mark-release-recapture (MRR) methodologies (Royle and Young 2008), and those that assign dispersing seeds to parent plants (Smouse and Sork 2004) or dispersing individuals to their origins (Berry et al. 2004; Schmidt et al. 2019). Approaches that summarise dispersal at the population level often aim to estimate the intergenerational dispersal parameter, σ, the standard deviation of parent-offspring separation distances (Wright 1943; Broquet and Petit 2009). Together with local population density (ρ), σ defines the neighbourhood size (N_w_ = 4πρσ^2^; Wright, 1946), the effective number of potential mates or parents within one generations’ dispersal range. While dispersal can be usefully assessed at both individual and population levels, recent studies in spatial population genetics have moved towards resolving this conceptual duality via frameworks that characterise dispersal at both levels (Aguillon et al. 2017; Bradburd and Ralph 2019).

The evolutionary consequences of dispersal can be understood in relation to dispersal barriers and dispersal ability. Barriers that restrict or redirect dispersal can have major effects on the distribution, connectivity and long-term persistence of wild populations (Ozinga et al. 2009; Berger-Tal and Saltz 2019). Recent and ongoing change in global land use has led to the addition of new barriers that are of particular concern for conservation (Debinski and Holt 2000) and the removal of pre-existing barriers that spread biological invasions along new pathways (Hulme et al. 2008). Studies investigating the impact of barriers that have recently changed or exhibit transient patterns of connectivity require methods capable of detecting their immediate effects (Landguth et al. 2010; Zeigler and Fagan 2014).

Besides barriers, dispersal ability itself works as an isolating force. Conspecific individuals distributed continuously through space are more likely to reproduce with nearby individuals when dispersal ability is limited relative to the spatial extent of the population, resulting in a decline in genetic similarity as distance increases (isolation by distance; Wright, 1943). Connectivity within and between populations is strongly influenced by σ, which determines the spatial scale at which isolation by distance operates (Broquet and Petit 2009). Both distance and barriers affect connectivity within populations, though determining their relative strength is often difficult when barriers at specific geographical positions also subdivide individuals spatially (Cushman et al. 2006). Unrecognised spatial structuring within populations can have strong effects on later inferences (Neel et al. 2013; Battey et al. 2020).

A recently developed genetic approach, Kindisperse (Jasper et al. 2019, 2021), estimates dispersal distance parameters from close kin data. Kindisperse estimates σ from the difference in variance between the spatial distributions of two or more kin categories, such as between full-sibs and full-first cousins. The specific kin categories used depends on the dispersal biology of the species, but individuals must typically be sampled at the same life stage (Jasper et al. 2021). In this regard, Kindisperse aims to summarise the dispersal variance across a single generation of the spatial pedigree (Bradburd and Ralph 2019), and can be considered a close-kin mark-recapture (CKMR) methodology (Bravington et al. 2016). This approach has operational advantages over MRR methods when applied to taxa that are difficult or costly to individually observe or where the ‘marking’ process is intrusive. Estimates of σ derived from close kin dyads have recently been validated *in silico* for different study designs (Jasper et al. 2021).

These same close kin data may also be useful for testing dispersal barriers. For instance, dyads of first order kin (parent-offspring or full-sib) observed on opposite sides of a geographical barrier indicate traversal of this barrier within the last generation or breeding cycle. As close kin provide information specific to the immediate past, they can assess barrier effects at very fine scales, such as where barriers are crossed in a single generation and individuals do not inhabit the barrier space. This may help evade lag-time issues associated with investigating gene flow immediately after a barrier is added or removed (Landguth et al. 2010), which makes close kin methods better suited to detecting recent change than methods that infer barriers from allele frequencies (Bradburd et al. 2013; Botta et al. 2015; Petkova et al. 2016). Several studies have explored kin-based barrier detection methods but these are by design either limited to assessing absolute barriers (Soanes et al. 2018; Schmidt et al. 2021) or they assess partial barriers but elide the effects of distance (Escoda et al. 2017, 2019); this effectively treats σ as being much larger than the distance between sampling points. A more versatile means of barrier investigation would be useful given that partial barriers are common and σ is often relatively small.

Here, we develop a method for using close kin dyads to detect dispersal barriers that accounts for the relative densities and distances between sampling points, and we apply it to a population of *Aedes aegypti* (yellow fever mosquito) sampled continuously through space in the urban environment of Jeddah, Saudi Arabia. Studies of *Aedes aegypti* dispersal frequently report the effects of barriers such as roads (Maciel-De-Freitas et al. 2007; Hemme et al. 2010; Schmidt et al. 2018; Regilme et al. 2021) but these have not estimated the strengths of specific barriers relative to other barriers and to distance. Here we do precisely that, and show how our method based on close kin performs favourably at this scale against methods that consider the genetics of the entire population. The same close kin data are also used to estimate σ with Kindisperse (Jasper et al. 2021), which has previously only been applied to individuals clustered within apartment buildings (Jasper et al. 2019). We also use isolation by distance patterns to estimate N_w_ (Rousset 2000). Given N_w_ = 4πρσ^2^, we then use N_w_ and σ to estimate ρ, the local population density.

Fine scale information about barriers, σ, N_w_, and ρ is critically important for this population of *Ae. aegypti,* as this study site is currently being assessed for mosquito releases aimed at establishing a transinfection of *w*AlbB *Wolbachia* in the local mosquito population following the success of releases with this strain in other areas (Nazni et al. 2019). *Wolbachia* are intracellular bacteria that inhibit dengue transmission when transinfected into *Ae. aegypti* (Walker et al. 2011), and *Wolbachia* can spread through wild mosquito populations and establish when transinfected mosquitoes are released in sufficient abundance (Barton and Turelli 2011; Schmidt et al. 2017). Establishment success can be strongly affected by barriers, which help maintain local infection frequencies above a critical frequency 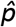 but also reduce the spread of *Wolbachia* into neighbouring regions (Barton and Turelli 2011; Schmidt et al. 2017). *Wolbachia* release outcomes are also influenced by σ, which determines the area over which mosquitoes should be released, and ρ, which determines how many mosquitoes should be released per unit area (Barton and Turelli 2011; Schmidt et al. 2017). As well as barriers and σ, this study provides a comparison of N_w_ in this and other *Aedes* populations, highlighting the utility of this parameter particularly when σ can be estimated. Finally, we compare dispersal estimates for *Ae. aegypti* derived from MRR studies with those derived from close kin dyads in this and one other study (Jasper et al. 2019).

## Materials and Methods

### Study site and sampling

*Aedes aegypti* were collected in Feb 2020 from an ~0.8 km^2^ urban area in Jeddah, Saudi Arabia (Figure 1A). This area covers part of the Al-Safa 9 region in the Almatar Municipality in Central Jeddah, where *Wolbachia-based* control operations are currently planned, as well as areas to the south and to the west of Al-Safa 9. Al-Safa 9 was separated from the southern area by a ~60 m wide road and the western area by a ~30 m wide road. Roads within this size range have been previously identified as potential dispersal barriers for *Ae. aegypti* (Russell et al. 2005; Schmidt et al. 2017). Most buildings in Al-Safa 9 are apartments of 3–6 storeys and pedestrian traffic is frequent throughout the region. *Aedes* mosquitoes are abundant year-round in Jeddah with a drop in numbers around December (Khan et al. 2018).

**Figure 1.**
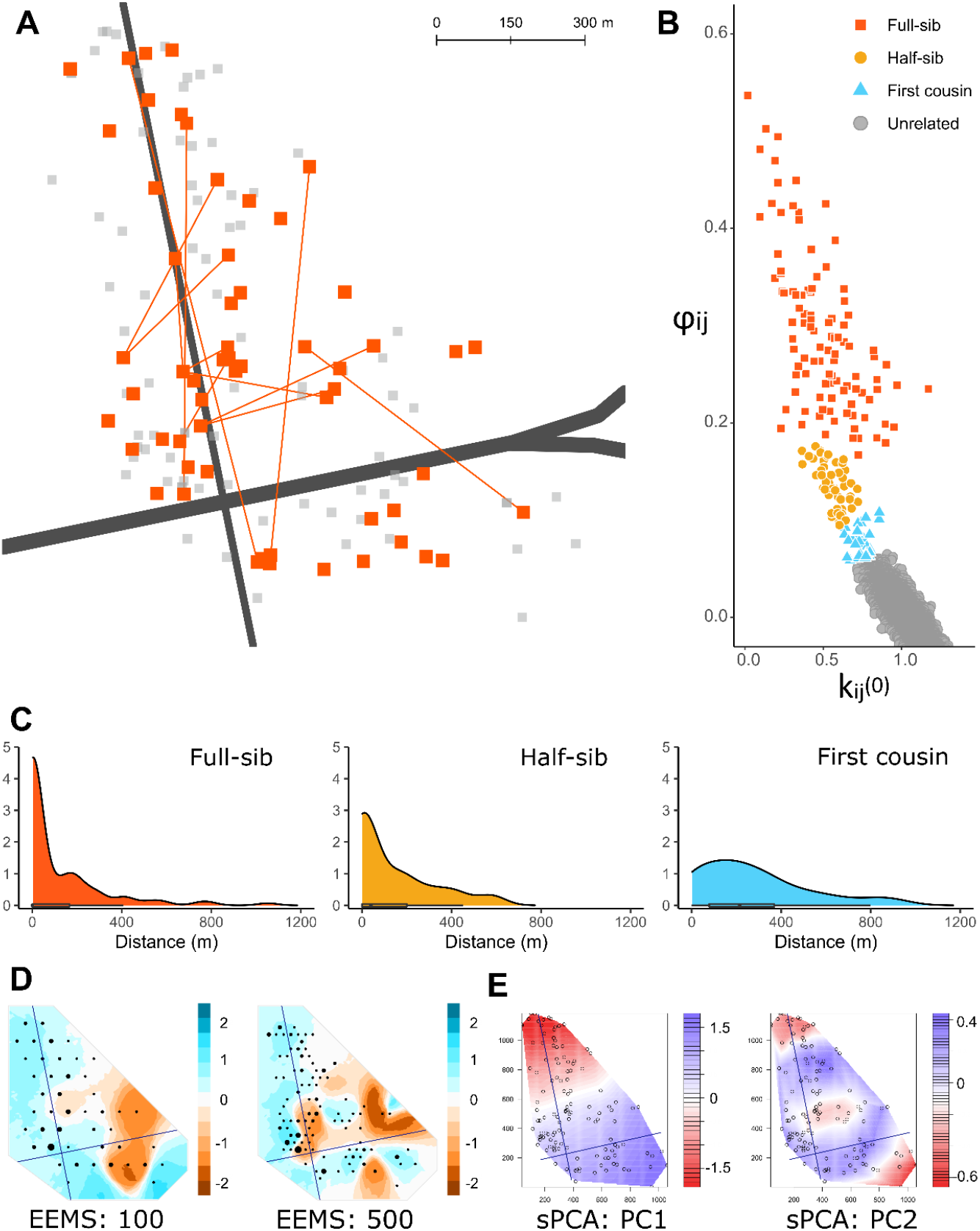
Traps, close kin dyads and spatial genetic structure in the Jeddah sample site. Figure 1A is a map of the study site, with grey squares indicating positive traps and large red squares indicating traps where full-sib dyads were found at different locations. Red lines show full-sib dyads found across the major roads, which are indicated in dark grey. Figure 1B shows the assignment of dyads to kinship categories based on kinship coefficient (φ_ij_) and probability of sharing zero alleles (k_ij_^(0)^). Figure 1C displays density plots of distances separating dyads of each kinship category. Figures 1D and 1E show results from EEMS (1D) and SPCA (1E), using one individual per trap and with road locations indicated. Figure 1D displays posterior mean migration rates, where blue indicates relatively higher migration and brown indicates lower migration, and where black circles show aggregations of positive traps assigned to grids of 100 or 500 demes. Figure 1E displays interpolation maps for the first two sPCA lagged scores, with similar colours indicating similar scores, and circles indicating locations of positive traps.

Six hundred ovitraps were deployed in the sampling area on Feb 3^rd^, 2020. Traps were individually georeferenced and deployed randomly through the sampling area, to collect *Ae. aegypti* across a continuous range of distances and on either side of each road. Each trap used a felt attached to a bucket containing water and 2-3 grass pellets to collect eggs from ovipositing *Ae. aegypti* females, which are laid on the felt. On Feb 11^th^, felts were collected and new felts deployed which were collected on Feb 18^th^. Of the 1200 felt samples taken over the two-week sampling period, 137 had *Ae. aegypti* eggs, 113 of which were collected in the second week. The 137 positive felts were taken from 128 unique locations (Figure 1A). Eggs collected from felts were hatched and reared to adulthood, and then stored in 100% EtOH at −20°C.

### Sequencing and genotyping

We generated double digest restriction-site associated DNA (ddRAD) sequence data from 266 *Ae. aegypti* (S1 Appendix), selected to maximise the spatial distribution of individuals. Protocols for DNA library preparation follow previous work (Rašić et al. 2014) but library size was restricted to 20 individuals to decrease variation in coverage within libraries (S1 Text). Libraries were sequenced using 150 bp paired-end chemistry at BerryGenomics Company (Beijing, China) on a NovaSeq 6000 (Illumina, California, USA).

We used process_radtags in Stacks v2.0 (Rochette et al. 2019) to demultiplex sequence reads, trim them to 140 bp, and discard reads with average Phred score below 20. We used bowtie v2.0 (Langmead and Salzberg 2012) to align reads to the *Ae. aegypti* genome assembly (Matthews et al. 2018) using --very-sensitive alignment. Individual single nucleotide polymorphism (SNP) genotypes were called via the Stacks v2.0 program ref_map. SNPs were retained when genotypes were called in ≥ 90% individuals and when ≥ 3 copies of the minor allele were detected (Linck and Battey 2019). We imputed missing genotypes in Beagle 4.0 (Browning and Browning 2016), then pruned SNPs by linkage disequilibrium (R package “SNPRelate” (Zheng et al., 2012); using the snpgdsLDpruning function with arguments, method = “r”, slide.max.n = 50 and ld.threshold = 0.005). This reduced the initial set of 81,780 SNPs to a set of 3,058 unlinked SNPs with no missing genotypes.

### Close kin identification

We used the R package ‘PC-Relate’ (Conomos et al. 2016) to estimate kinship and identity by descent parameters for all 35,245 possible dyads. PC-Relate controls for genetic structure such as from isolation by distance by conditioning the data with principal components (PCs). Two principal components (PCs) were used to condition for genetic structure as no structure was observed in additional PCs (S1 Figure). Parameters included the kinship coefficient (φ∪) and the probabilities of sharing zero (k_ij_^(0)^) or two (k_ij_^(2)^) alleles at a site, and these were used to determine each dyad’s order of kinship. As all samples were collected within a two-week period and represent the same generation, we assumed that 1^st^ order dyads were full-sibs, 2^nd^ order dyads were half-sibs, and 3^rd^ order dyads were first cousins. Previous work has found that at fine scales almost all isolation by distance is caused by such close kin dyads rather than ‘background’ structure (Aguillon et al. 2017).

Dyads of 1^st^, 2^nd^, and 3^rd^ order kinship have expected φ_ij_ of 0.25, 0.125, and 0.0625 respectively. Accordingly, dyads with φ_ij_ > 0.1875 are most likely 1^st^ order kin, 0.1875 > φ_ij_ > 0.0938 are most likely 2^nd^ order kin, and 0.0938 > φ_ij_ > 0.0469 are most likely 3^rd^ order kin. However, more accurate assignments can be obtained using both φ_ij_ and k_ij_^(0)^ (Figure 1B). Unrelated, first cousin, and half-sib dyads are expected to align along a line intersecting the x-axis at k_ij_^(0)^ = 1 and the y-axis at φ_ij_ = 0.25, but where full-sibs have k_ij_^(0)^ scores ~0.25 higher (Conomos et al. 2016). Using both parameters to assign dyads resolved several instances of impossible pedigrees (e.g. three individuals containing two full-sib dyads and one half-sib dyad) that would have arisen if only φ_ij_ was used.

Assigning 3^rd^ order dyads was difficult even when considering φ_ij_ and k_ij_^(0)^ together (Figure 1B). We reasoned that if unrelated dyads were incorrectly classed as first cousins this could bias dispersal estimates upwards. This is because distances between cousins represent two generations of dispersal while distances between unrelated dyads represent additional generations. However, if first cousins were incorrectly classed as unrelated, this would simply reduce the sample size rather than introduce biases, as the Kindisperse method for *Ae. aegypti* considers only sibs and first cousins (Jasper et al. 2021). Considering the uncertainty around estimates, we assigned first cousin dyads conservatively and treated the ‘first cousin’ category as a composite category containing both full and half-first cousins (Jasper et al. 2019). Relatedness parameters and location data for all dyads are in S1 Appendix.

### Road barrier testing

#### i) Frequency of full-sib dyads across roads

We tested whether the western or southern roads were functioning as dispersal barriers using the spatial distribution of trap pairs containing full-sib dyads (Figure 1A). In this study, each individual was sampled as an egg, and thus the distance between any full-sib dyad represents the movement of only a single, unsampled individual – their mother. When two full-sibs in a dyad are observed on either side of a putative barrier, this implies their mother crossed this barrier while moving between ovitraps to oviposit. If a given feature is genuinely a barrier to movement, then fewer full-sib dyads should be observed on opposite sides of the barrier than on the same side, relative to expectations from the number of dyads and the distances between traps.

To test for barriers, we first considered three treatments: trap pairs on the same side of the road; trap pairs across the western road; and trap pairs across the southern road. Considering first the trap pairs on the same side of the road, 30 of these contained one or more pairs of full-sibs, and these had a maximum separation of 510 m. There were 2442 trap pairs on the same side of the road within 510 m. We used these values to divide the distance of 0 – 510 m into 10 distance classes, with each set to contain an equal number of trap pairs on the same side of the road (i.e. 244 – 245) (S1 Table). For each of the ten distance classes, we calculated how many pairs of traps contained full-sibs (n_full-sib traps_). We used the ratio of 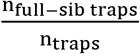 to produce expected rates of full-sib detection at each distance class (where n_traps_ = 244 – 245), which we treated as expected rates when barriers are not present. As no full-sibs were observed beyond 510 m, we could not calculate ratios for these distances.

We compared these expected rates of detection against rates of detection observed for the western and southern road treatments, which each had a putative barrier. These used the same 10 distance classes, but n_traps_ was different for each treatment. For instance, for the first distance class, 0 – 70.8 m, there were 244 traps on the same side of the road, 14 across the western road, and 2 across the southern road. For each distance class, we used these n_traps_ scores and the expected ratios of full-sibs to produce expectations of n_full-sib traps_, rounding to the nearest integer (S1 Table). Differences between expected and observed n_full-sib traps_ at each distance class were compared using Wilcoxon signed-rank tests for paired samples.

We compared the performance of this full-sib approach against three commonly employed methods for detecting barriers: distance-based redundancy analysis in the r package ‘vegan’ (dbRDA: Legendre & Anderson (1999)), spatial principal components analysis (sPCA) in the R package ‘adegenet’ (Jombart 2008) and estimating effective migration surfaces (eems: Petkova, Novembre, & Stephens (2016)). Details of these analyses are in S2 Text.

#### ii) Separation distances of full-sib dyads across roads

While fewer full-sib dyads are expected across barriers, these dyads are also predicted to be separated by greater distances than when no barrier is operating. This is because longer-range dispersal (e.g. > 500 m) in *Ae. aegypti* tends to be anthropogenically associated, such as via transportation in cars (Eritja et al. 2017). These passive movements will not be affected by barriers in the same way as active flight, and thus long-range movement may not be reduced in frequency despite barriers restricting short-range movement.

To test this prediction, we calculated for each of the three road treatments (same side, across the western road, across the southern road) the ratio of n_full-sib traps_ found within the *i*th percentile of trap distances relative to the total nfull-sib traps for that treatment, with *i* = {10%, 20%, … 90%} (S2 Table). Thus, 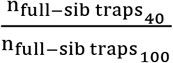 indicates the proportion of trap pairs containing full-sibs that are found within the nearest 40% of trap distances. Differences in ratios between treatments were assessed with an ANOVA.

### Dispersal parameterisation

#### i) Estimating σ from close kin dyads

We estimated dispersal location kernels (Nathan et al. 2012) using the Kindisperse approach (Jasper et al. 2019, 2021). Kindisperse uses variance addition and subtraction to isolate specific dispersal location kernels from the spatial distributions of close kin dyads. We estimated two kernels. The ovipositional dispersal kernel (σ_ovi_) describes the spatial distribution of eggs as a result of females moving between ovitraps to oviposit. This movement of ovipositing females is called ‘skip’ oviposition when it occurs within one gonotrophic cycle, as in this study (Christophers 1960). The parent-offspring dispersal kernel (σ or σ_PO_) describes an entire generation of dispersal, life stage to life stage (i.e. egg(t-1) to egg(t)). For all analyses, we retained only a single dyad of a given order found in the same trap or trap pair, to avoid bias from sequencing unequal numbers of individuals across traps.

The spatial distribution of full-sibs was used to estimate σ_ovi_. We estimated σ_ovi_ with the Kindisperse function ‘axpermute’, using a vector of full-sib distances as input and setting ‘composite = 2’ as each full-sib dyad represented two draws from this kernel (i.e. σ^2^_Fs_ = 2σ^2^_ovi_). It is important to clarify that, as our samples were collected as eggs, and as we included full-sib dyads from the same traps, the σ_ovi_ kernel is not intended to summarise female movement distances *per se.* Rather, it represents the distribution of related individuals at a specific life stage (i.e. egg) within a specific generation as a result of female movement (Jasper et al. 2021). The same σ_ovi_ kernel can also be estimated from full-sibs at the adult life stage that have moved from their locations at the egg stage, though confidence intervals will be broader (Jasper et al. 2021).

The σ_PO_ kernel was estimated from the difference in variances between the spatial distribution of sibs and cousins. This analysis considers the recent coancestry of close kin dyads, where sibs coalesce in the parental generation and cousins coalesce in the grandparental generation. For *Ae. aegypti,* if the spatial distribution of a full-sib dyad reflects two draws from the mother’s σ_ovi_ kernel, the spatial distribution of a full-first cousin dyad will reflect two draws from the grandmother’s σ_ovi_ kernel and two draws from the parent’s dispersal kernel from egg through to oviposition (i.e. σ^2^_1C_ = 2σ^2^_ovi_ + 2σ^2^_PO_) (Jasper et al. 2021). Using the additive properties of variance, the difference between the full-sib and full-first cousin dispersal kernels will be equal to two ‘draws’ from the σ_PO_ kernel (i.e. σ^2^_1C_ - σ^2^_FS_ = 2σ^2^_PO_). If, as in this dataset, the first cousin category contains full and half-first cousins, σ_PO_ is estimated by compositing the full-sib category with the half-sib category (Jasper et al. 2019, 2021).

We estimated σ_PO_ with the ‘axpermute_standard’ function, using vectors of full-sib and first cousin distances as input, and using ‘amixcat = “H1C”’ and a vector of half-sib distances to ‘composite’ the full-sib category (‘bcompcat = “HS”’).

#### ii) Estimating N_w_ from isolation by distance

As well as the above close-kin based analyses, we estimated neighbourhood size (N_w_) using the inverse of the slope of the regression of linear genetic distance among individuals against ln-transformed geographical distance and omitting dyads within σ_PO_ (Rousset 2000). We compared these results against N_w_ estimates from other *Aedes* populations. These included *Ae. aegypti* populations from urban high-rise apartments in Kuala Lumpur, Malaysia (N = 162) (Jasper et al. 2019) and urban Cairns, Australia (N = 171) (Schmidt et al. 2018) and an *Ae. albopictus* population from islands in northern Australia (N = 301) (Schmidt et al. 2021). All sequence data from the *Ae. aegypti* populations were processed and filtered identically to the Jeddah samples. As the Cairns sample was taken shortly after *Wolbachia* releases in the area, we estimated N_w_ separately for pairs with *Wolbachia* (N = 59) and pairs without *Wolbachia* (N = 112); there was no isolation by distance pattern between pairs of different infection status. The *Ae. albopictus* estimates omitted pairs found on different islands. For the Cairns and *Ae. albopictus* N_w_ calculations we assumed σ_PO_ = 96.5 m.gen^-1/2^, the average of the estimates for Al-Safa and Malaysia (46.0 m.gen^-1/2^).

## Results

### Close kin identification

We identified 190 full-sib dyads, 64 half-sib dyads, and 46 first cousin dyads from the 35,245 dyads (Figure 1B). Most (104) full-sib dyads were from the same trap. All but 2 full-sib dyads were sampled in the same week, indicating likely movement within a single gonotrophic cycle. The was no relationship between the distance separating full-sib dyads and either φ_ij_ (R^2^ = 0.005) or k_ij_^(0)^ (R^2^ = 0.006). Maximum distances among close kin were 1052 m (full-sib), 588 m (half-sib) and 890 m (first cousin) (Figure 1C).

### Road barrier testing

#### i) Frequency of full-sib dyads across roads

There were 44 trap pairs containing full-sib dyads. Thirty were on the same side of the road, while 12 were across the western road and 3 across the southern road, including a single trap pair across both roads (Figure 1A, Figure 2A, S1 Table). Within the 10 distances classes that ranged 0 – 510 m, there were 9 trap pairs with full-sib dyads across the western road and 0 across the southern road (Figure 2B). These observed values are lower than expected values for each treatment, with 19 trap pairs with full-sib dyads expected across the western road (Wilcoxon test, z = 1.221, P = 0.111) and 9 expected across the southern road (Wilcoxon test, z = 2.222, P = 0.013). The 8^th^ distance class (365–429 m) was omitted from calculations as it did not contain any full-sibs for any treatment which was likely a stochastic effect.

**Figure 2.**
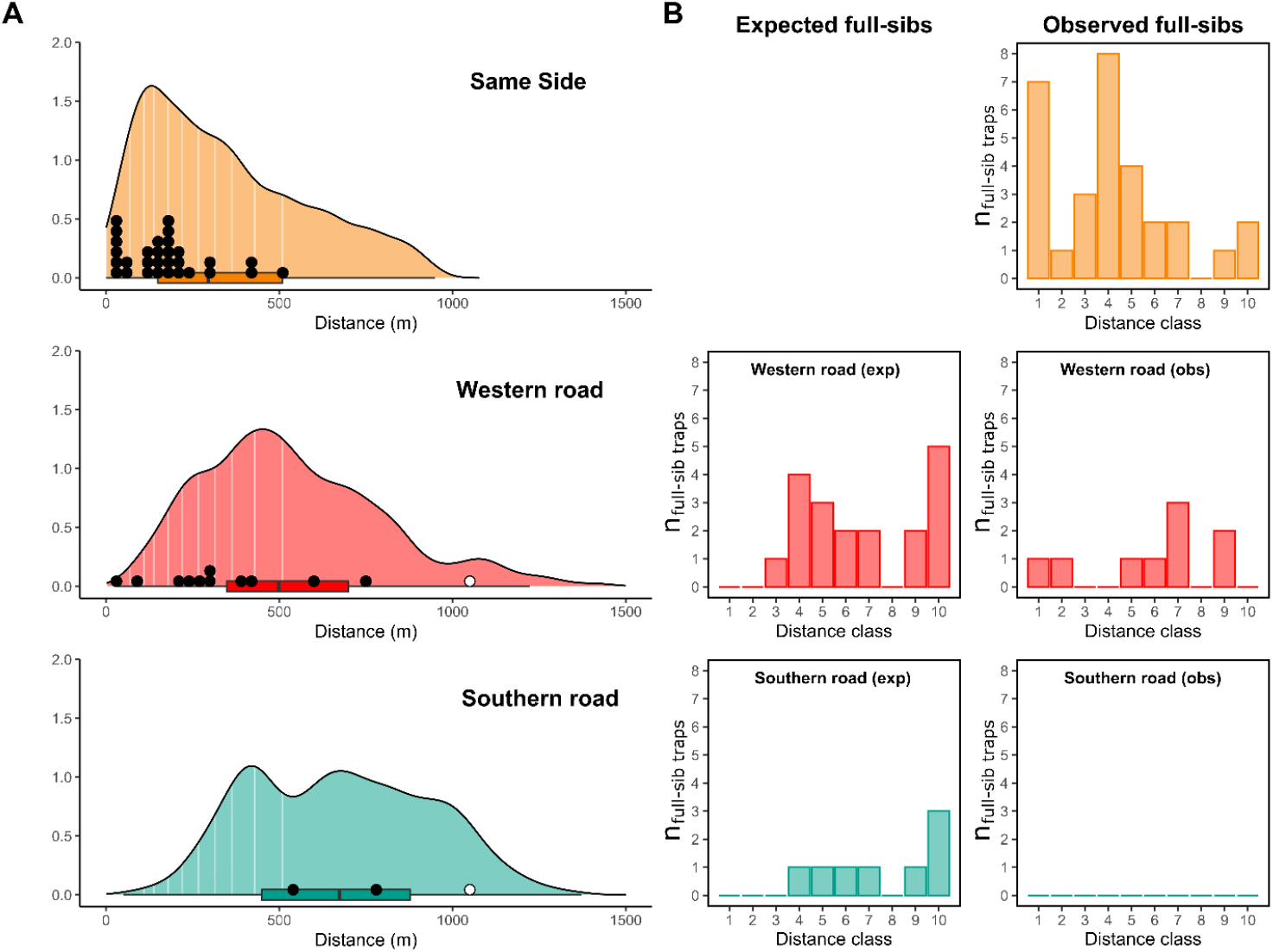
Testing road barrier effects. The density plots in Figure 2A describe the pairwise distances between traps for each of the three road treatments. The black circles indicate the distances between each pair of traps containing full-sibs, and the white circles indicate a single full-sib pair found across both roads. The 10 distance classes are delineated by the vertical line segments through each plot. In Figure 2B, the rate of full-sib detection on the same side of the road (yellow, top right) is used to generate expected distributions of full-sibs across each road (exp) which are compared against observed distributions (obs).

None of the dbRDAs found either of the road barriers to have a statistically significant effect, regardless of whether analyses considered the whole dataset or tested each road individually, or whether geographical variables were used to condition the model. However, higher F values and lower P values were observed for the southern road barrier (whole dataset: F_1,135_ = 1.233, P = 0.089) than for the western road barrier (whole dataset: F_1,135_ = 0.982, P = 0.525), a similar pattern to the close kin dyads. Both EEMS treatments (100 deme and 500 deme) indicated a region of lower migration around the southern road barrier though the exact barrier location was difficult to pinpoint from this (Figure 1D). The first lagged score (PC1) of the sPCA showed an isolation by distance pattern between north and south but no specific structure around the southern road barrier (Figure 1E).

#### ii) Separation distances of full-sib dyads across roads

Traps with full-sib dyads across the southern road were separated by relatively larger distances than those on the same side or across the western road (F_2,24_ = 5.62, P = 0.0099). For the same side, western, and southern road treatments, the 50^th^ percentile of trap distances contained 1624, 2115, and 1643 traps within distances of 296 m, 499 m, and 648 m, respectively. The proportions of full-sibs within these distances were 90%, 75%, and 33%, respectively (S2 Table).

### Dispersal parameterisation

#### i) Estimating σ from close kin dyads

Dispersal parameterisation used 202 trap pairs: 105 full-sibs 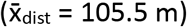, 49 half-sibs 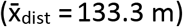, and 41 first cousins 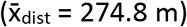. The spatial distribution of full-sib dyads indicated that the ovipositional dispersal location kernel (σ_ovi_) was 108.3 m.gen^-1/2^ (95% C.I. 76.4–139.7) (Table 1). The spatial distributions of the sib and cousin dyads indicated that the parent-offspring dispersal location kernel (σ_PO_) was 146.9 m.gen^-1/2^ (95% C.I. 81.5–197.1). As dyads across the southern road barrier had different spatial distributions, we also ran analyses with these omitted, giving σ_ovi_ = 84.1 m.gen^-1/2^ (95% C.I. 61.5–105.4) and σ_PO_ = 142.3 m.gen^-1/2^ (95% C.I. 78.2–193.4). Dispersal estimates are given in m.gen^-1/2^ as we are modelling dispersal in two dimensions, where σ is an axial standard deviation of a bivariate distribution (Wright 1946).

**Table 1.**
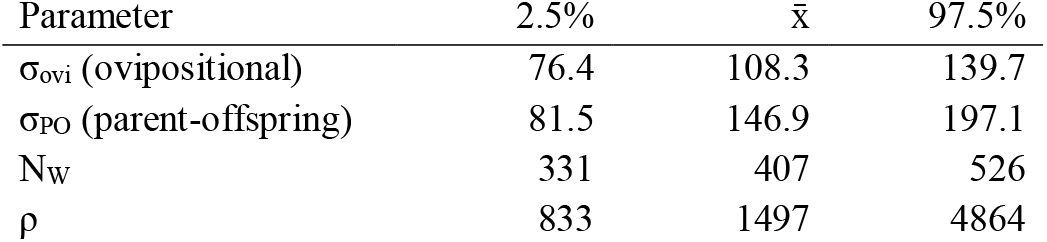
Population parameter estimates for Jeddah *Ae. aegypti.* Dispersal estimates are in m.gen^-1/2^, ρ is in km^-2^.

| Parameter | 2.5% | $\bar{x}$ | 97.5% |
| --- | --- | --- | --- |
| $\sigma_{\text{ovi}}$ (ovipositional) | 76.4 | 108.3 | 139.7 |
| $\sigma_{\text{PO}}$ (parent-offspring) | 81.5 | 146.9 | 197.1 |
| $N_W$ | 331 | 407 | 526 |
| $\rho$ | 833 | 1497 | 4864 |

We compared these *Ae. aegypti* dispersal estimates with estimates derived from mark-release-recapture (MRR) studies (Figure 3; S3 Table). These studies do not comprise an exhaustive review of the MRR literature as in Guerra et al. (2014); here we focus on more recent research. For each study reporting dispersal estimates from multiple releases we only include two estimates, selected randomly. Estimates are compared against two parameters of study design that vary among dispersal studies: the area over which sampling was conducted (Figure 3A) and the number of specific sampling points per unit area (Figure 3B), both of which were square root transformed to enforce proportionality with dispersal estimates. MRR estimates are compared against close kin dyad estimates from this study and from Jasper et al. (2019), which have been converted from σ_PO_ into mean Euclidean distances (r) expected under Gaussian assumptions using: σ^2^ = 0.5 × E[r^2^] (Crawford 1984; Rousset 2004).

**Figure 3.**
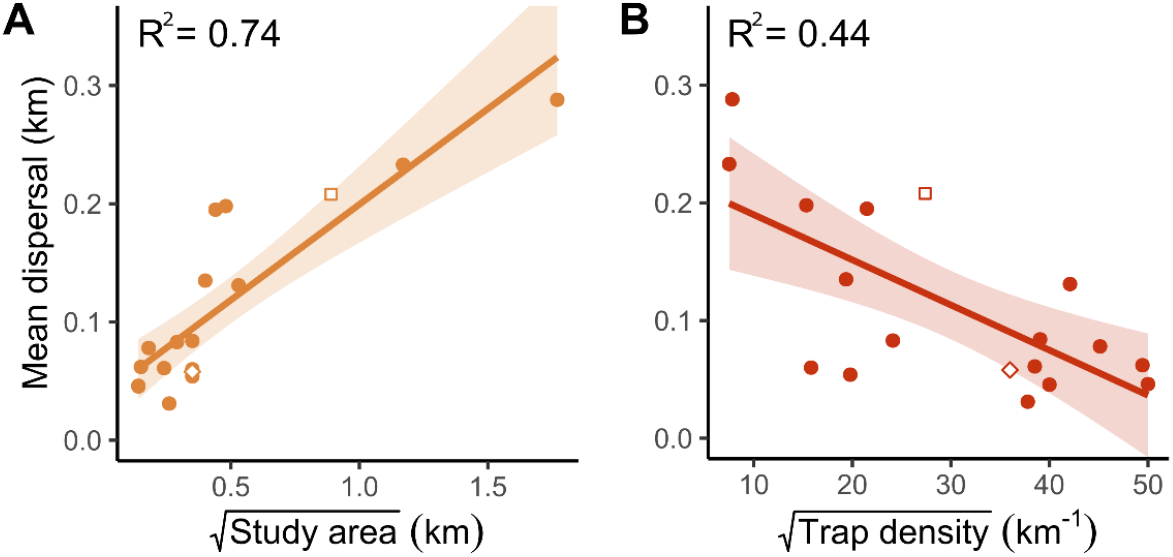
Comparison of Euclidean dispersal distance estimates across studies. Estimates from MRR are indicated by solid filled circles. Estimates from close kin dyads are a white square (this study) and a white lozenge (Jasper et al. 2019). Data and references for each estimate are listed in S3 Table. (Muir and Kay 1998; Ordóñez González et al. 2001; Harrington et al. 2005; Russell et al. 2005; Maciel-De-Freitas et al. 2007; Maciel-De-Freitas et al. 2007; Maciel-de-Freitas and Lourenço-de-Oliveira 2009; Lacroix et al. 2012; Valerio et al. 2012; Winskill et al. 2015; Marcantonio et al. 2019; Juarez et al. 2020; Trewin et al. 2021).

Dispersal studies conducted over larger areas have generally reported larger estimates of dispersal (Figure 3A, R^2^ = 0.74). This is expected as long-distance movement will only be detected in studies conducted across sufficient area, a pattern noted previously in empirical (Guerra et al. 2014) and simulated (Jasper et al. 2021) investigations. However, studies conducted across large areas tend to also sample from a more dispersed set of points, as it is unfeasible to maintain a high density of traps across large areas. This lower density of trapping correlates with larger estimates of dispersal (Figure 3B, R^2^ = 0.44). As *Ae. aegypti* often disperse over very short distances and stay within a single building their entire life (Harrington et al. 2005; Jasper et al. 2019), study designs with traps spaced far from each other and from release points may fail to record this short-range movement and thus produce inflated estimates.

#### ii) Estimating N_w_ from isolation by distance

After omitting dyads within the axial dispersal distance of σ_PO_ (146.9 m.gen^-1/2^), neighbourhood size (N_w_) in Al-Safa was estimated at 407 (95% C.I. 331 – 526). Using the equation, N_w_ = 4π × σ_PO_^2^ × ρ, where ρ is the effective density of *Ae. aegypti,* we estimate ρ = 1497 km^-2^ (95% C.I. 833 – 4864). Figure 4 shows how the estimates of N_w_, σ_PO_ and ρ from Al-Safa compare with estimates from other *Aedes* populations. N_w_ was relatively consistent among the *Aedes* populations considering the ~2-3 orders of magnitude breadth in N_w_ recorded among and even within species (Battey et al. 2020). Likewise, estimates were consistent for the *Wolbachia*-infected (N_w_ = 944) and uninfected (N_w_ = 1016) subsamples from Cairns, Australia. While *Ae. aegypti* in Al-Safa had a larger N_w_ than those from Kuala Lumpur (407 vs 124), σ_PO_ was also larger at Al-Safa 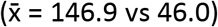, leading to a lower estimated ρ than in Kuala Lumpur (1497 vs 4,663).

**Figure 4.**
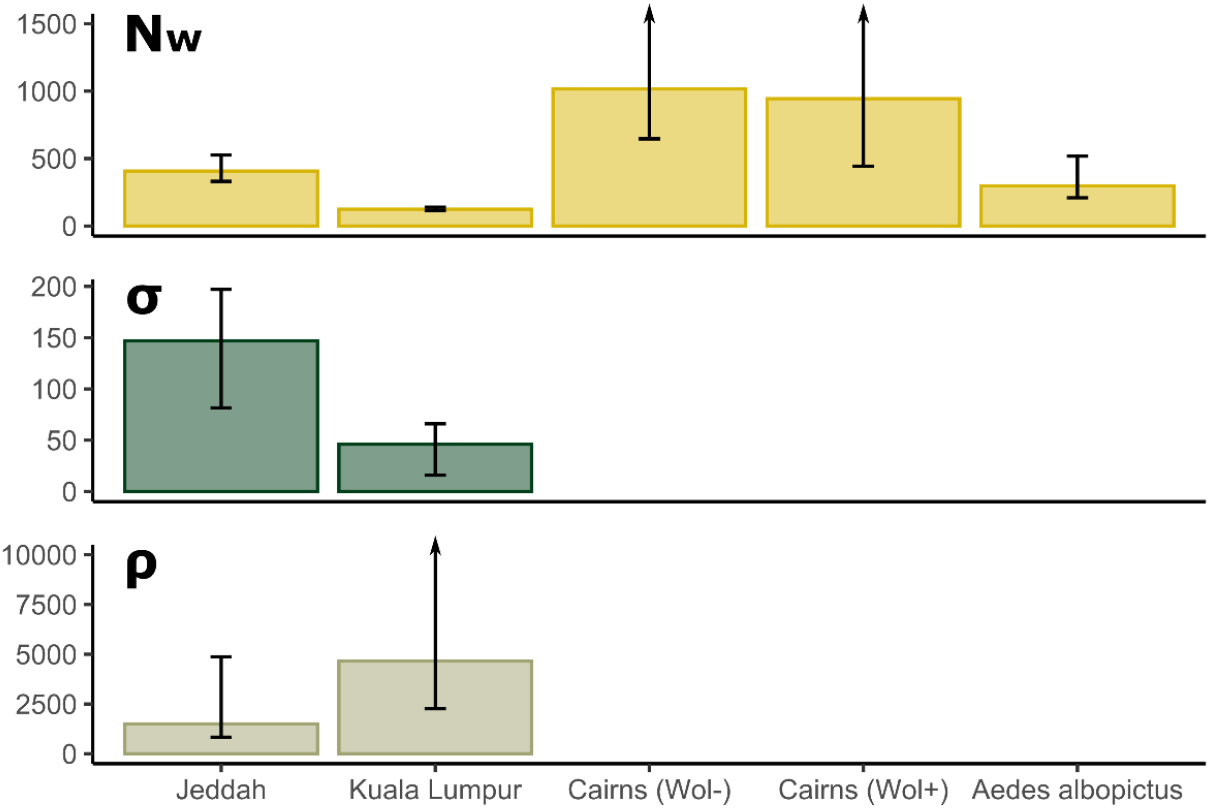
Neighbourhood size (N_w_), dispersal (σ_PO_), and density (ρ) estimates for *Aedes* populations. Units for σ are m.gen^-1/2^, units for ρ are km^-2^. Unplotted 95% C.I. estimates are: Cairns (Wol-) N_w_ = 2374; Cairns (Wol+) N_w_ = ∞; Kuala Lumpur ρ = 38,550.

## Discussion

Dispersal shapes the fate of many wild populations and is a frequent focus of ecological and evolutionary research. A conceptual duality in early studies in whether dispersal should be investigated as a life-history event (Howard 1960) or an intergenerational process (Wright 1943) has led to a corresponding duality in approach that continues to the present day (Broquet & Petit, 2009). Current genetic approaches that can detect dispersal across discrete generations are helping to resolve this duality (Bradburd and Ralph 2019; Jasper et al. 2021). This study has used close kin dyads sampled from the same life stage and generation to reveal the specific movement of individuals and to estimate average dispersal patterns within the study site over recent generations. The locations of full-sib dyads were used to infer the specific movements of mothers, and these movement patterns revealed fine-scale dispersal barriers with greater sensitivity than methods assessing overall genetic differentiation. The difference in variance between the sib and cousin spatial distributions provided an estimate of σ, and spatial genetic differentiation over multiple generations indicated N_w_ and ρ. This framework thus uses a single dataset to investigate dispersal as both a life-history event and an intergenerational process.

While σ is often treated as constant across all populations of a species (Nunney 2016), σ may vary when a species like *Ae. aegypti* inhabits a broad range of environments. Our estimate of σ for Al-Safa *Ae. aegypti* (146.9 m.gen^-1/2^) was much higher than that of a population inhabiting Malaysian high-rise apartments (46.0 m.gen^-1/2^, Jasper et al. 2019), and could reflect differences in behaviour or physiology. However, in the Malaysian population, *Ae. aegypti* are clustered within apartment buildings and do not inhabit the spaces between them. Lower estimates of σ are thus expected given that mosquito movement is limited to mostly short distances (within apartments) with occasional long distance movement (between apartment buildings) and little intermediate movement. If a continuously-distributed population like Al-Safa was sampled in a similarly clustered manner, this absence of intermediate distances would downwardly-bias estimates. Nevertheless, the lower σ in the Malaysian high-rise apartment buildings remains an accurate summary of the intergenerational dispersion of alleles, though it may yet be underestimated due to sampling over too small an area (Figure 3A; Guerra et al. (2014); Jasper et al. (2021)). Estimates from Malaysia and Al-Safa should thus be cautiously compared.

N_w_ is known to vary among and even within populations (Shirk and Cushman 2014; Battey et al. 2020). This may reflect variation in local σ as well as density (ρ). The interpretation of N_w_ is sometimes confused with N_e_ (effective population size) (Nunney 2016); while N_w_ is limited to describing local spatial structure it can be usefully applied to continuously-distributed populations where σ^2^ is much smaller than area covered by the population (Leblois et al. 2003), whereas in continuous populations Ne can be subject to strong biases from sampling (Neel et al. 2013). The *Aedes* N_w_ estimates reported here (124-1016) describe variation among populations relating directly to ρ and σ, and which can be compared with N_w_ estimates from Anopheline mosquitoes (~150-700, Clarkson et al. 2020) and *Drosophila pseudoobscura* (~570-1100, Dobzhansky & Wright 1943). By comparison, N_e_ estimates in *Ae. aegypti* alone exhibit much greater variance. Resampling methodologies give estimates of 25–3,000 (Saarman et al. 2017) and coalescent methods give contemporary estimates in post-bottleneck invasive-range populations from ~5000 in Mexico (Crawford et al. 2017) to ~1,500,000 across the Caribbean (Sherpa et al. 2018). The difference between these latter estimates may reflect the spatial scales over which individuals were aggregated for analysis, which does not affect N_w_ in the same way (Neel *et al.* 2013, c.f. Clarkson et al. 2020).

Reconciling individual-based movement observations such as those obtained from MRR with dispersal estimates from molecular data is often difficult (Broquet & Petit, 2009). However, here we find our estimates of Euclidean distance derived from σ accord well with estimates from MRR studies conducted at similar scales (Figure 3). In general, choice of dispersal inference method will depend on the study system. Close kin methods are particularly suitable for short-lived organisms for which dispersal is spatially limited and difficult to observe directly. Close kin can also reveal occasional dispersal over long distances (Escoda et al. 2017; Schmidt et al. 2021). By contrast, MRR methods are particularly effective when applied to larger animals observed at multiple timepoints such as through camera traps (Silver et al. 2004; Royle and Young 2008). An additional advantage of close kin methods is that density (ρ) can be estimated from N_w_ estimates when σ is known (Rousset 2000; Leblois et al. 2003). Running this operation in reverse, in which ρ is used to estimate σ, is also viable (Broquet et al. 2006), but requires density data which can be difficult to collect.

Fine-scale estimation of barriers, σ, N_w_, and ρ can inform management strategies for wild populations, particularly those experiencing recent change. For *Ae. aegypti* in the Al-Safa 9 region, these will inform *Wolbachia* releases. The major strategic implication of these findings is that the release site is bordered by barriers, with the southern barrier (60 m width) stronger than the western barrier (30 m width). Accordingly, if the invasion is to either spread outwards or retreat inwards from Al-Safa (c.f. Schmidt et al. (2017)) this will more likely occur at the weaker western barrier than at the southern barrier. Other undetected barriers within the study site may also have these effects, but the effects of the two roads are the most critical as they define the region within which releases will take place. Considering σ, the much larger estimate of σ = 146.9 m.gen^-1/2^ at Al-Safa suggests dispersal distances are higher than in Malaysia even if Malaysian estimates are slightly downward-biased by small study site dimensions (Jasper et al. 2021). With higher dispersal in Al-Safa, we would expect to observe faster dynamics of spread and reinvasion, and *Wolbachia* will need to be deployed over a relatively larger area than in Malaysia (Barton and Turelli 2011). Estimated mosquito population densities were also much lower at Al-Safa (~1500 km^-2^) than in the Malaysian site (~4500 km^-2^). Estimated densities are also likely lower in Al-Safa than in Cairns. This is because N_w_ in Al-Safa (407) was much lower than in Cairns (944-1016), and the rate of *Wolbachia* spread through Cairns was sufficiently slow such that σ > 100 m.gen^-1/2^ is unlikely (Schmidt et al. 2017). Thus, the lower N_w_ in Al-Safa probably reflects a lower relative density than Cairns. This Cairns sample was taken from around the Parramatta Park region where trap collections were consistently high for this city and *Wolbachia* frequencies were relatively stable after establishment (Schmidt et al. 2017). On the other hand, *Wolbachia* frequencies were more volatile at the lower-density Edge Hill release site in Cairns (Schmidt et al. 2017), which might also occur at Al-Safa, making it important to monitor *Wolbachia* frequencies carefully.

### Conclusions

Ongoing change in global land use brings with it the addition and removal of dispersal barriers, much of which has occurred recently relative to generation time (Zeigler and Fagan 2014). Following previous work showing how close kin dyads can estimate σ (Jasper et al. 2021), this paper has demonstrated their use in barrier detection. Close kin methodologies can identify how barriers affect different forms of dispersal such as active flight and passive movement when these operate over different spatial scales, and should be particularly useful for detecting recent changes to barriers down to the scale of a single generation or reproductive cycle. Even so, we have shown that close kin methods can outperform methods that consider the genetics of the entire population in assessing barriers that have existed for 100s of generations, provided the investigation is conducted at a sufficiently fine scale. This strong signal at fine scales accords with close kin findings in other taxa (Aguillon et al. 2017). Importantly for barrier detection, only data from first-order kin is required, allowing the use of cheaper genetic markers such as microsatellites that can capture first-order relationships (Hauser et al. 2021).

---

**S1 Text: DNA extraction, library preparation and sequencing**

DNA was first extracted using DNA kits, either Qiagen DNeasy Blood & Tissue Kits (Qiagen, Hilden, Germany) or Roche High Pure™ PCR Template Preparation Kits (Roche Molecular Systems, Inc., Pleasanton, CA, USA).

To build ddRAD libraries, we digested 100 ng of DNA from each individual in a 45 μL volume, consisting of 9 units each of MluCI and NlaIII restriction enzymes (New England Biolabs, Beverly MA, USA), NEB CutSmart buffer, and water. Digestions were run for 3 hours at 37 °C with no heat kill step, and the products were cleaned with 1.5× SpeedBead Magnetic carboxylate (Cytiva, USA). These were ligated to Illumina P1 and P2 adapters (Peterson *et al.* 2012) overnight at 16 °C with 1,000 units of T4 ligase (New England Biolabs, Beverly, MA, USA), followed by a 10-minute heat-deactivation step at 65 °C. We performed size selection with a Pippin-Prep 2% gel cassette (Sage Sciences, Beverly, MA) to retain DNA fragments of 300–450 bp.

Final libraries were amplified by eight PCR, each of which used 1 μL of size-selected DNA, 5 μL of Phusion High Fidelity 2× Master mix (New England Biolabs, Beverly MA, USA) and 2 μL of 10 μM standard Illumina P1 and P2 primers. These were run for 12 PCR cycles, then cleaned and concentrated using 0.8× paramagnetic beads. Each ddRAD library contained 20 mosquitoes, and were sequenced using 150 bp paired-end chemistry at BerryGenomics Company (Beijing, China) on a NovaSeq 6000 (Illumina, California, USA).

**S2 Text: Methods for dbRDA, sPCA and EEMS**

For the three analyses, we subsampled a single random individual from each felt (N = 137) to avoid including full-sibs from the same oviposition.

We ran separate dbRDAs to test each barrier, using a binary variable to encode the barrier and a distance matrix of Rousset’s *a* scores as the dependent variable, calculated in SPAGeDi (Hardy & Vekemans 2002). We ran multiple dbRDAs for each road (R package ‘vegan’, function *capscale),* first without any geographical distance variable, then with a distance variable coded as the entire set of principle coordinates produced from a matrix of geographical distances (function *pcnm),* then with only those principal coordinates that had P < 0.05 when assessed against Rousset’s *a* scores directly. We also reran these dbRDAs on reduced datasets that omitted traps from across the road not being analysed, providing direct comparisons between the central zone and the western (N = 106) and southern (N = 90) regions. We also tried including all individuals (N = 266) but this led to strong geographic biases from traps containing multiple close kin.

We ran sPCA using a ‘neighbourhood by distance’ connection network, and plotted the interpolated scores of the first two principal components to observe whether geographical genetic structure could be linked to the positions of the roads. For eems, we used default input parameters but set the number of demes at either 100 or 500 for comparison.

**S1 Table:**
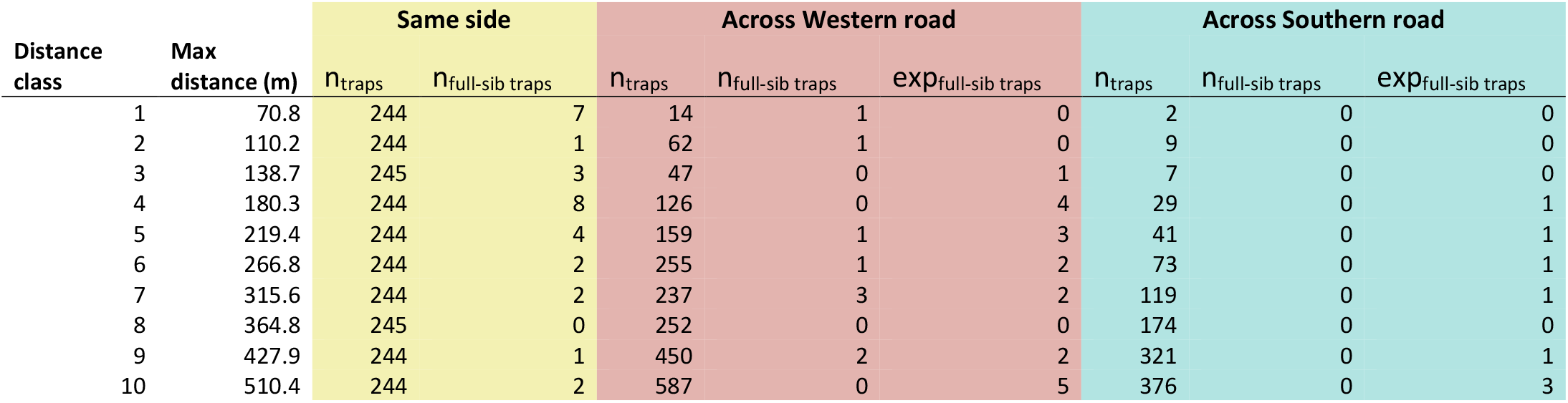
Trap and full-sib data for assessing road barriers I: observed and expected values at the ten distance classes.

**S2 Table:** Trap and full-sib data for assessing road barriers II: ratios of full-sib traps within each percentile.

| Percentile | Same side |  | Across Western road |  | Across Southern road |  |
| --- | --- | --- | --- | --- | --- | --- |
|  | Trap distance<br>(m) | ratio<sub>full-sib traps</sub> | Trap distance<br>(m) | ratio<sub>full-sib traps</sub> | Trap distance<br>(m) | ratio<sub>full-sib traps</sub> |
| 10 | 84 | 0.27 | 224 | 0.27 | 333 | 0.00 |
| 20 | 128 | 0.37 | 305 | 0.58 | 410 | 0.00 |
| 30 | 179 | 0.63 | 381 | 0.58 | 469 | 0.00 |
| 40 | 232 | 0.83 | 440 | 0.75 | 552 | 0.00 |
| 50 | 296 | 0.90 | 499 | 0.75 | 648 | 0.33 |
| 60 | 364 | 0.90 | 567 | 0.75 | 734 | 0.33 |
| 70 | 452 | 0.97 | 653 | 0.83 | 825 | 0.67 |
| 80 | 571 | 1.00 | 747 | 0.83 | 940 | 0.67 |
| 90 | 711 | 1.00 | 879 | 0.92 | 1067 | 1.00 |

**S3 Table:**
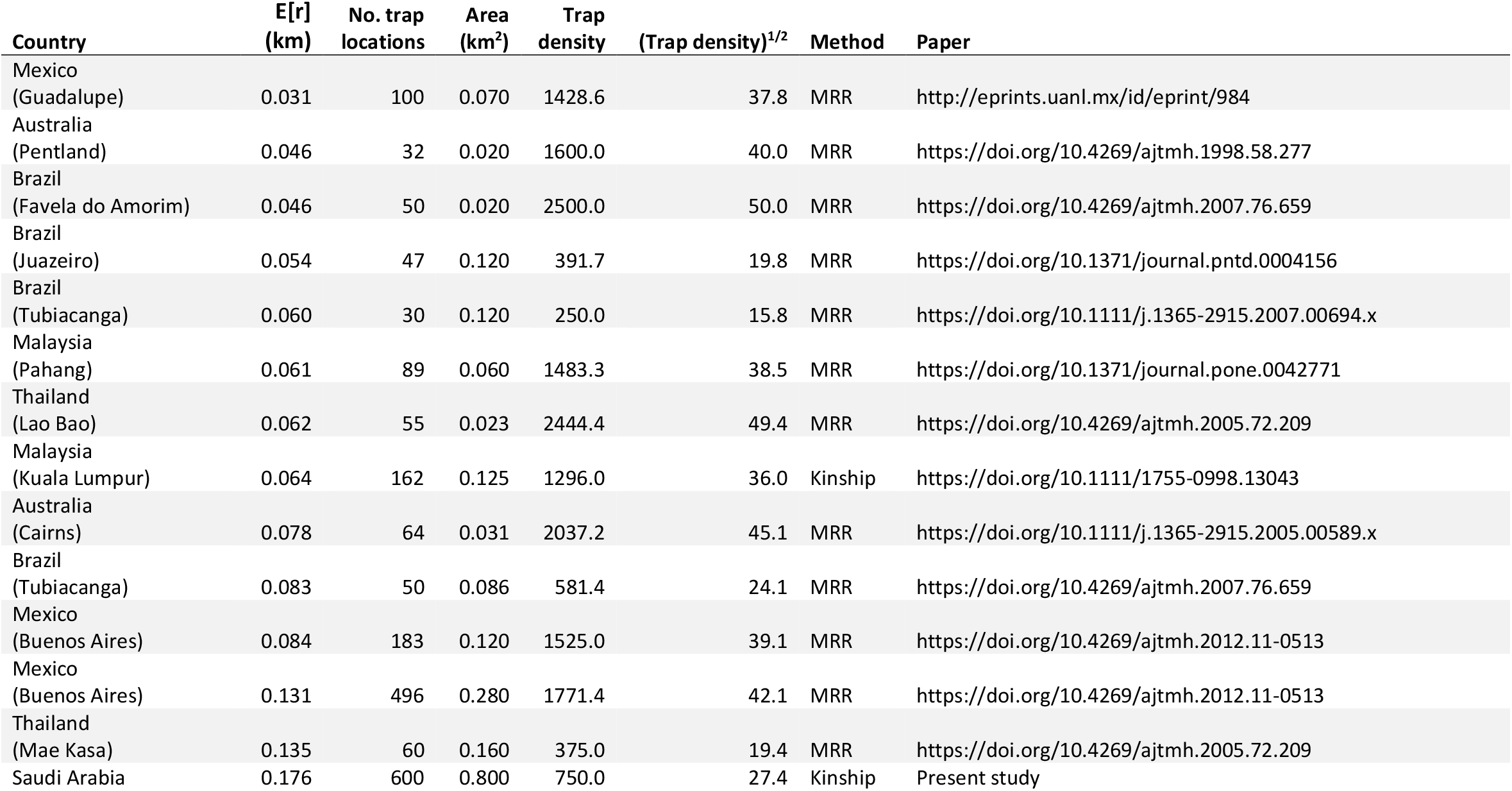
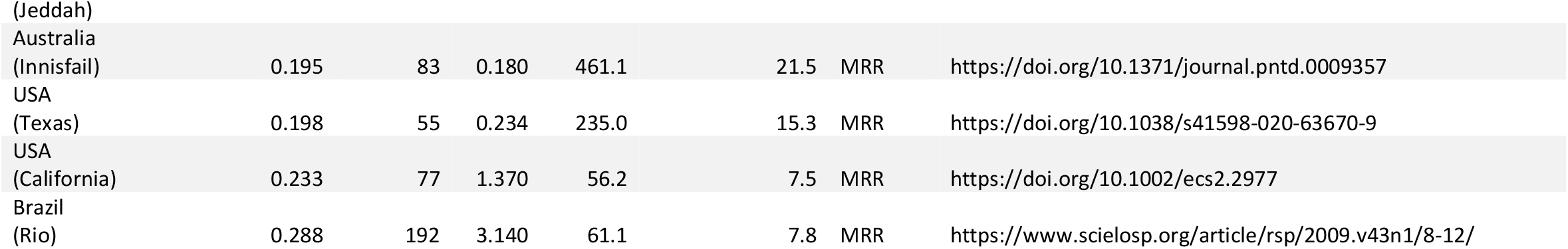
Data used in comparing dispersal estimates across *Aedes aegypti* studies.

**S1 Figure.**
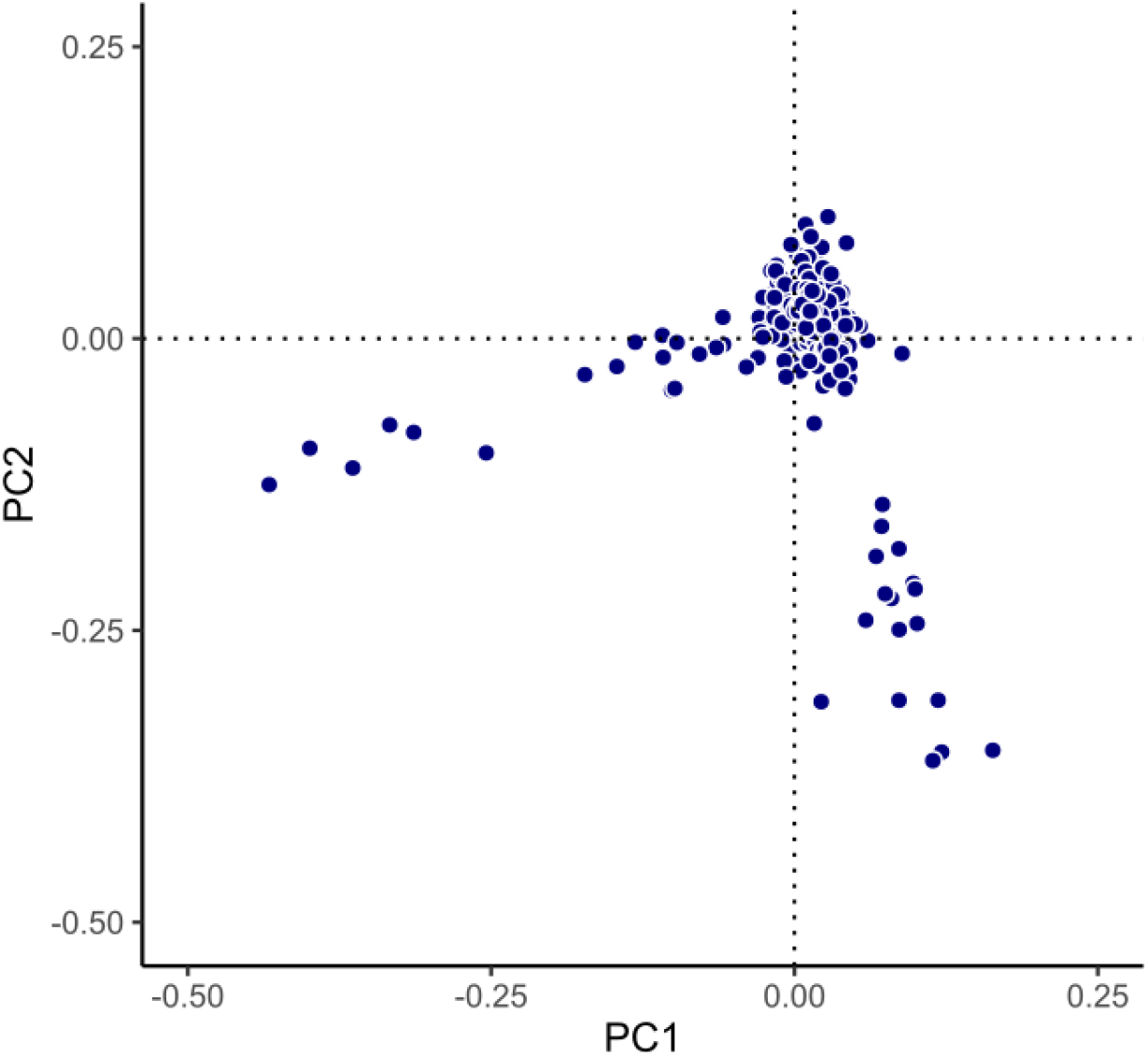
First and second principal components (PCs) used in PC-Relate.

